# Nanopore sequencing provides rapid and reliable insight into microbial profiles of Intensive Care Units

**DOI:** 10.1101/2021.05.14.444165

**Authors:** Guilherme Marcelino Viana de Siqueira, Felipe Marcelo Pereira-dos-Santos, Rafael Silva-Rocha, María-Eugenia Guazzaroni

**Affiliations:** Faculdade de Filosofia Ciências e Letras de Ribeirão Preto (FFCLRP-USP), Ribeirão Preto, SP, Brasil. 14040-901; Faculdade de Medicina de Ribeirão Preto (FMRP-USP), Ribeirão Preto, SP, Brasil. 14049-900

**Keywords:** nanopore sequencing, Illumina sequencing, 16S rRNA, environmental monitoring, intensive care units, healthcare-associated infections

## Abstract

Fast and accurate identification of pathogens is an essential task in healthcare settings. Next generation sequencing platforms such as Illumina have greatly expanded the capacity with which different organisms can be detected in hospital samples, and third-generation nanopore-driven sequencing devices such as Oxford Nanopore’s minION have recently emerged as ideal sequencing platforms for routine healthcare surveillance due to their long-read capacity and high portability. Despite its great potential, protocols and analysis pipelines for nanopore sequencing are still being extensively validated. In this work, we assess the ability of nanopore sequencing to provide reliable community profiles based on 16S rRNA sequencing in comparison to traditional Illumina platforms using samples collected from Intensive Care Units from a hospital in Brazil. While our results point that lower throughputs may be a shortcoming of the method in more complex samples, we show that the use of single-use Flongle flowcells in nanopore sequencing runs can provide insightful information on the community composition in healthcare settings.

## Introduction

Surveillance and control of organisms commonly found in the microbiome of surfaces surrounding patients in hospital settings are among the main public health challenges faced worldwide. Healthcare-associated infections (HAIs) are one of the leading causes of patient morbidity and mortality and are often associated with prolonged hospitalizations that weigh on health systems (1,2). Moreover, HAI-related organisms have been shown to persist in environments and are strongly correlated with antimicrobial resistance (AMR), which imposes difficulties in treatment and contributes to the emergence of new multidrug resistant pathogens in healthcare settings (2).

Massive parallel sequencing technologies are important tools to aid monitoring of healthcare environments (3). While traditional pathogen detection methods usually involve laborious cultivation and biochemical assays, molecular characterization of organisms using specific biomarkers, such as the 16S rRNA gene, bypasses these steps, enabling the identification of poorly-represented or even unculturable pathogens at a much faster rate (3,4).

The minION (Oxford Nanopore Technologies - ONT) is one of the latest installments of high-throughput third-generation sequencers available. This nanopore-driven device was introduced to the market in the mid 2010s, and quickly gained popularity as a tool for clinical and environmental monitoring and (meta)genomic profiling due to its capacity to generate extremely long reads while remaining highly portable and relatively cheaper than most standard sequencing technologies (5,6). Long reads are of particular importance for microbial community profiling, given that the 16S rRNA gene cannot be entirely sequenced by traditional second-generation platforms and the choice of which variable region(s) to amplify directly affects which genera can be detected with these machines (7). Moreover, the often time-consuming library preparation steps and longer sequencing runs hold back the widespread adoption of second-generation devices for routine applications in healthcare settings, where rapid results are critical (8).

Despite gaining traction, nanopore sequencing protocols and analysis pipelines are still being validated by the community at large, and many research groups have been thoroughly evaluating its performance against traditional (usually Illumina) platforms. In the present study, we investigated how nanopore-based long-read sequencing compares to Illumina sequencing for the study of complex microbial communities from hospital surfaces. In order to identify potentialities and limitations of nanopore sequencing we resequenced a set of samples collected in a previous study that used Illumina’s MiSeq to explore the microbial diversity within Intensive care Units in a Brazilian hospital (9). Despite the difference in approaches, the results obtained in this study are comparable to the findings reported in the original publication and support the same conclusions. Thus, nanopore sequencing may be implemented as a fast and accurate methodology for tracking the distribution of harmful microbial species throughout healthcare facilities.

## Material and methods

### Samples acquisition

The samples used in this study were collected by a collaborator in a hospital in the city of Ribeirão Preto in the state of São Paulo (Brazil) in 2018. The procedures used to acquire the samples are fully described in a previous publication of our group (9). In short, selected surfaces from Intensive Care Units (ICUs) and Neonatal Intensive Care Units (NICUs) were thoroughly streaked with sterile swabs premoistened with sterile Amies media for two minutes, after which the swabs were placed in sterile 15 mL Falcon tubes containing an additional 1mL of sterile Amies media and kept under refrigeration until DNA extraction was performed. Metagenomic DNA was extracted using the MoBio Powersoil DNA isolation kit, and samples were stored in a −80°C freezer until further processing (9).

### Library preparation

In the present study, approximately 5 ng of these samples were used as template for barcoding PCR using primers provided in ONT SQK-16S024 sequencing kit. PCR amplifications were made with Phusion polymerase (NewEngland Biolabs) under the standard reaction protocol, including the use of DMSO 3%. After an initial denaturation step at 98°C for 2’30” the reaction cycles in the PCR program were set as follows: denaturation at 98°C for 15”, annealing at 52°C for 15”, extension at 72°C for 01’30”, with a final extension step of 72°C for 5’ after the 35^th^ cycle.

Success of 16S amplification was assessed in 0.8% agarose gels stained with SYBR-Safe^™^ (Thermo Fisher Scientific). In order to avoid loss of material, we did not perform a purification step before pooling the amplicons together. Instead, samples that showed well-defined ≈1500 bp bands were mixed at proportional volumes, starting with 5 μL for the strongest bands. The pools were then purified with AMPure XP beads (Beckman Coulter) and resuspended in 10 mM Tris-HCl pH 8.0 with 50 mM NaCl as recommended by ONT. Nanodrop^™^ One (Thermo Fisher Scientific) was used to assess concentration of the purified pools. If needed, concentrations were adjusted to 10 - 20 ng/μL. Finally, 5 μL of each pool were mixed with 0.5 μL of the Rapid adapter (RAP) of the SQK-16S024 sequencing kit and incubated for 5 minutes at room temperature.

Library preparation, as well as Flongle flowcells priming and loading steps were then carried according to instructions described in the manufacturer’s protocol (version: 16S_9086_v1_revI_14Aug2019). Sequencing runs lasted for up to 24h and were performed in MinION model Mk1B (ONT).

### Long-read data processing

FASTQ files were generated and demultiplexed concurrently to sequencing using Guppy basecaller (version 4.0.9) in the fast basecalling setting. After the runs ended, passed reads had barcode sequences removed using guppy_barcoder command line utility (ONT). NanoFilt (version 2.7.1) (10) was used to filter reads based on quality (QScore > 10) and size (1350 - 1650 bp), and suitable reads were finally aligned to the NCBI refseq 16S database using Minimap2 (version 2.17) (11). The alignment output was parsed in R (version 4.0.5) with the pafr package (version 0.0.2) (12). Only unique alignments with overlaps > 1000 bp were considered for further analysis.

A GitHub repository containing the scripts that we have developed for automatization of our pipeline can be found at https://github.com/GuazzaroniLab/nanopore_ICU_profiling

### Comparative data analysis

In our previous work, the V4 region of the 16S rRNA gene was amplified from metagenomic DNA samples and sequenced in 2×300 bp Illumina MiSeq runs. QIIME version 1.9.1 was used to determine Operational Taxonomic Units (OTUs) after sequencing (9). Data from the previous publication was retrieved and used for comparison with nanopore-generated results. Nanopore and Illumina datasets were compared in R (version 4.0.5) using vegan (version 2.5.7) (14) and the Tidyverse suite of packages (version 1.3.1) (15). For the nanopore dataset, bacterial taxa assigned to less than five reads in each sample were not considered in the analysis. Determination of microbial biomarkers in ICU wards was performed using LEfSe command-line tool version 1.0 (16)

## Results

### Nanopore sequencing can paint a reliable picture of the community composition of (N)ICUs despite its limited throughput

In a previous work from our group, samples from inanimate surfaces of (N)ICUs of a hospital in Brazil were collected and sequenced using an Illumina platform (MiSeq) after amplification of the ≈300 bp V4 region of the 16S ribosomal RNA gene (9). As shown by the diagram in **Figure 1**, in the present study, these samples were resequenced using minION, a nanopore-based platform, which allows full-length sequencing of the 16S gene.

**Figure 1.**
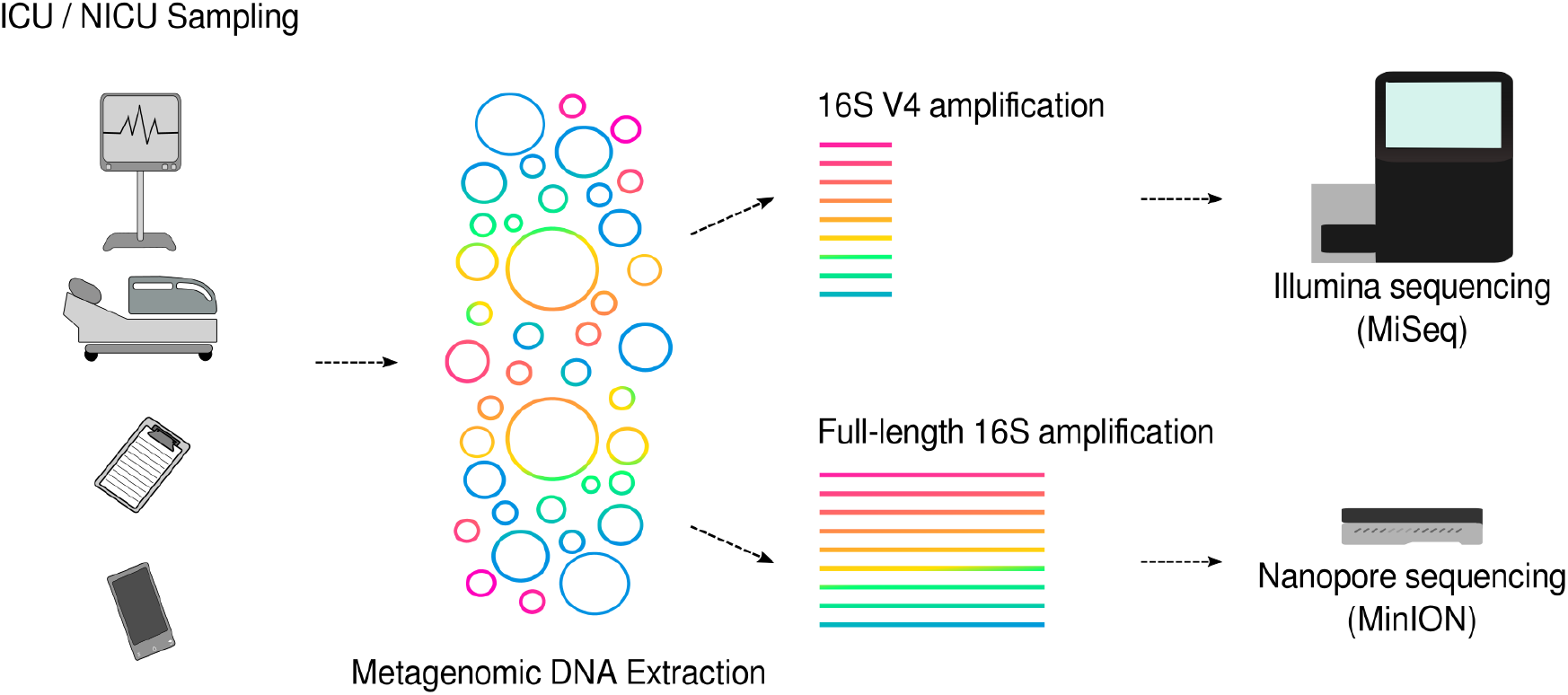
Overview of our experimental design. In a previous work of our group. metagenomic DNA was extracted from surfaces and objects surrounding patients in (N)ICUs of a hospital in Brazil. Samples were then subjected to V4 16S sequencing using Illumina MiSeq for microbial community profiling. In the present study, full-length 16S nanopore sequencing was performed in a subset of these samples, allowing us to compare both methods. For consistency, sample names followed the same code used in our previous work: sample source (e.g. “Handles” for door handles, “Ventilators”, “Mobiles” for staff cell phone devices, “Monitors” for cardiac monitors…), followed by either ICU or NICU and the ward (a, b or ab). For concurrent cleaning analysis, an uppercase “A” identifies samples collected after the cleaning. A full description of the samples can be found in our previous publication (9).

Our results seem to indicate that throughput in nanopore sequencing runs was limited in comparison to the amount of data previously generated by Illumina. This observation might be related to the use of Flonge flowcells in our sequencing runs. Introduced to the market in 2019 as a cheaper alternative to the standard conventional minION flowcell for smaller sequencing experiments, single-use Flonge flowcells have up to 126 nanopores available for sequencing instead of the standard 2048 present in the regular flowcell, and provides theoretical yields of up to 2.8 Gbp, according to the manufacturer. In the two runs that comprised this work there were, respectively, 83 and 60 pores available for sequencing at the start of the experiments. The minION sequencing runs generated 503 Mbp and 542 Mbp in passed reads. The number of passed reads ranged between 1611 (sample Ventilator-ICUaA) and 34083 reads (sample Pump-ICUa), averaging at 14296 reads across our samples **(Table S1)**. For comparison, in our previous work, 4.94 Gbp were generated by Illumina’s MiSeq platform, with an average read count per sample of 34621.

In addition to this, our strict processing pipeline resulted in a massive removal of data. On average, 80% of reads did not meet our criteria of i) having QScore > 10 (i.e., 90% of basecalling accuracy) and ii) being as large as 1650 bp but not smaller than 1350 bp. Aside from having the highest number of raw reads, sample Pump-ICUa also had the highest loss (92.5%) of reads after processing.

As a direct consequence of the differences in throughput between sequencing platforms, estimates of diversity within samples in the Illumina dataset are generally greater than those of the nanopore **(Figure S1)**. Nonetheless, hierarchical clustering of the samples shows a high level of agreement at genus level between the two approaches, with correlation indexes ranging from 60.3% to 95.6% in different subsets of our samples **(Figure S2)**. Moreover, rarefaction curves show that the vast majority of the nanopore-sequenced samples had a number of counts that far exceeded the requirement for saturation even after processing **(Figure S3)**. These results seem to indicate that, even with a lower throughput, nanopore sequencing with Flongle flowcells was able to provide satisfactory amounts of data and has the potential to support subsequent community analysis.

### Both sequencing approaches revealed shortcomings of the cleaning protocols employed to sanitize ICU surfaces

A main finding in our previous work was the apparent unevenness of efficacy of the concurrent cleaning procedures employed in the ICU wards. In fact, community composition analysis not only showed that the most abundant genera present in the samples collected before the cleaning also comprised the majority of the microbiota after the sanitation procedures took place, but that there was also a noticeable increase in the relative abundance of certain genera in the surfaces after the cleaning (9).

These conclusions can also be reached when assessing our nanopore sequencing dataset. Despite the differences in throughput and sensitivity between the two approaches, there is a great similarity between the most abundant bacterial genera detected in samples both with Illumina and nanopore sequencing, as shown in **Figure 2A**. In the five sites with comparative cleaning information that were resequenced, *Bacillus* was the predominant genus before cleaning (on average, 52% of relative abundance estimated with nanopore and 34.9% with Illumina), followed by genera like *Staphylococcus* (11% with nanopore versus 10.6% with Illumina), *Stenotrophomonas* (4.36% with nanopore versus 5.6% with Illumina), *Pseudomonas* (6.8% with nanopore versus 7% with Illumina), *Pseudoxanthomonas* (4,75% with nanopore versus 5,49% with Illumina) and *Castellaniella* (8.1% with nanopore versus 4.35% with Illumina). Due to the phylogenetic similarity between the genera *Escherichia* and *Shigella,* Illumina pipelines often report their relative abundances as *Escherichia/Shigella,* but our direct alignment approach actually reports each genus independently, even though their sequences might be too similar for accurate discerning based on 16S alone (17). Before cleaning, analysis of nanopore data shows that *Shigella* was present with relative abundance of 5.70%, while *Escherichia* accounted for 5.65%, while Illumina reported *Escherichia/Shigella* at 6.12% relative abundance.

**Figure 2.**
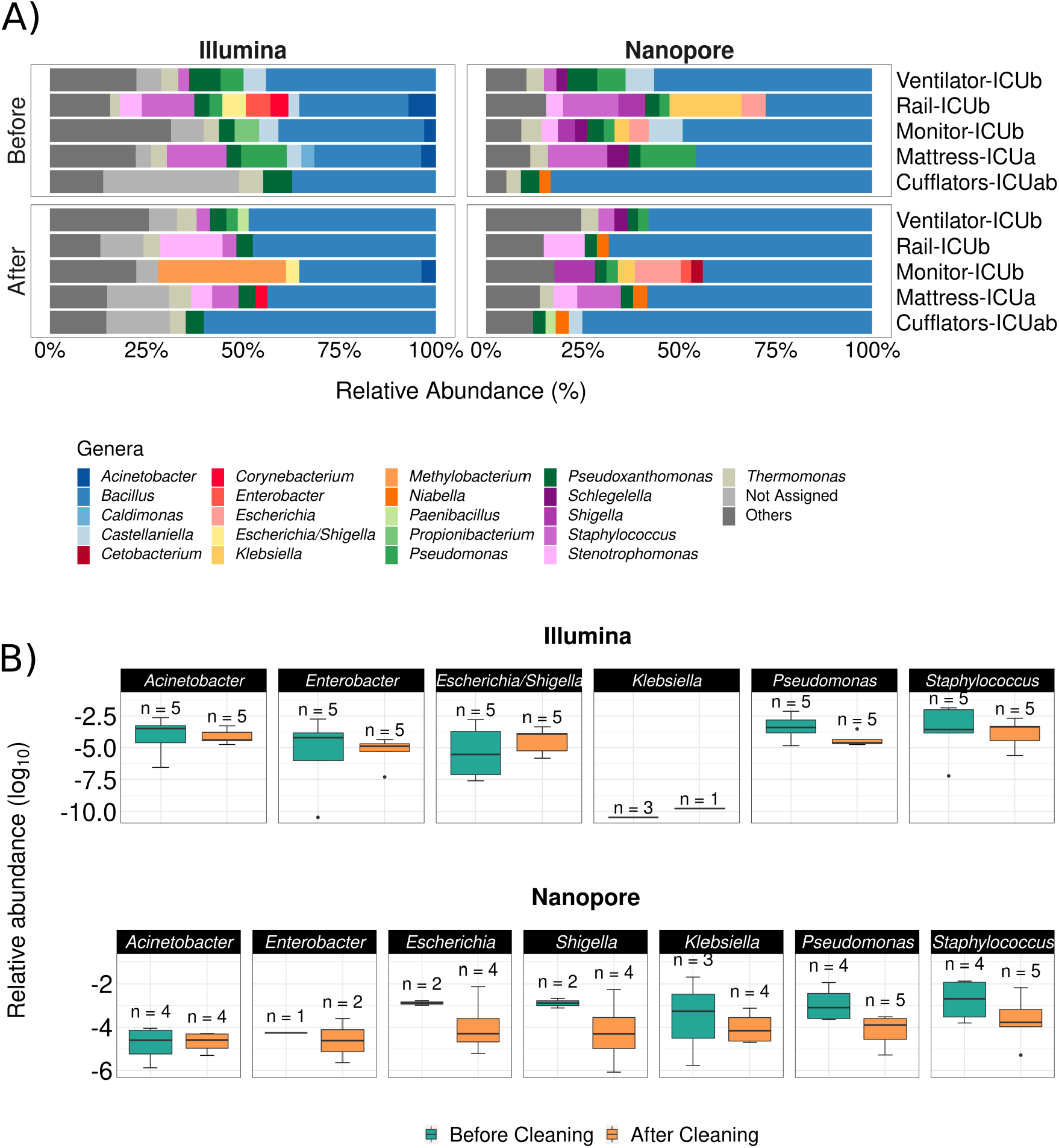
Comparison of the effect of cleaning routines in the ICU microbial profiles according to Illumina and nanopore sequencing. **A)** Stacked bar plots depicting main genera detected in ICU samples collected before and after concurrent cleaning with both approaches. Genera with relative abundances below 2.5% were grouped under the label “Others”. **B)** Boxplots show the effect of sanitation procedures on the abundance of selected HAI-related organisms detected in *n* different samples.

Considering 2.5% of relative abundance as a cutoff to determine the most abundant genera in the samples, the genera *Klebsiella*, *Schlegelella* and *Niabella* were not detected in the Illumina dataset while *Enterobacter*, *Proprionibacterium*, *Acinetobacter*, *Corynebacterium* and *Caldimonas* were absent from the nanopore dataset.

After cleaning, *Bacillus* remained the predominant genus in the samples (60% with nanopore versus 46.30% with Illumina), followed by *Stenotrophomonas* (8,41% with nanopore versus 10.8% with Illumina), *Staphylococcus* (7.7% with nanopore versus 4.6% with Illumina), *Pseudomonas* (2.8% with nanopore versus 2.94% with Illumina) and *Pseudoxanthomonas* (3% with nanopore versus 4.4% with Illumina). Moreover, in addition to the three genera described above, *Castellaniella*, *Cetobacterium* and *Enterobacter* were not detected by Illumina after cleaning, whereas *Methylobacterium*, *Acinetobacter* and *Corynebacterium* were absent from the nanopore dataset. After cleaning, *Escherichia/Shigella* relative abundance dropped in the Illumina dataset, reaching 3.48.%. However, the opposite was reported for genera *Escherichia* and *Shigella* when analyzed with the nanopore approach, with relative abundances of respectively 11.9% and 10.5% in the samples they were present.

Despite the general agreement between the two datasets, we notice that the prevalence of key genera associated with hospital infections across different ICU sites before and after cleaning varies greatly with the chosen method. As shown by **Figure 2B**, while it is clear that Illumina sequencing was able to detect the majority of investigated genera in all five sites, even if at lower relative abundances, there is a remarkable absence of *Klebsiella,* detected only at relative abundances of ≈10^-10^ in few samples. With nanopore, on the other hand, we retrieved all of the genera in fewer sites and were able to detect *Klebsiella* at greater relative abundances before and after cleaning. It should be noted that *Klebsiella pneumoniae* was the most abundant species (23,2%) from a total of 108 bacterial strains - distributed among 12 genera - that were detected by standard cultivation methods and Vitek 2 identification system in biological samples of hospitalized patients at the same period of the time of our sampling.

Overall, nanopore-generated estimates of the presence of pathogens in different samples were more conservative than those provided by Illumina. However, both approaches hint towards a lack of efficacy of the concurrent cleaning methods employed by hospital staff. Our results support that these practices do not seem to be adequate to sufficiently remove several HAI-related genera from inanimate surfaces and can end up serving as potential routes of cross-contamination of pathogens across hospital sites.

### Site-specific taxonomic biomarkers detected by Illumina and nanopore platforms allow spatial monitoring of the microbiota in ICU wards

In addition to exploring the differences in microbial profiles as a result of the cleaning methods employed by ICU staff, the investigation of differences in the composition of the microbiota across sections of the hospital in our previous study could explain the prevalence of certain nosocomial diseases in these sites (9).

In consonance with the results presented in the previous section, there was a high level of agreement between nanopore and Illumina regarding the most abundant genera in the NICU samples. At a cutoff level of 2.5% in relative abundance, there were 32 genera detected with Illumina and 28 with nanopore. The shared genera include common HAI-related organisms such as *Serratia*, *Bacillus*, *Delftia*, *Haemophilus*, *Stenotrophomonas*, *Acinetobacter*, *Streptococcus* and *Staphylococcus*. Genera like *Aliterella*, *Pseudopropionibacterium*, *Alloiococcus*, *Klebsiella*, *Kingella*, *Cylindrospermum*, *Massilia*, *Enterococcus* and *Aggregatibacter* were exclusive to the nanopore dataset, whereas *Propionibacterium*, *Marinomonas*, *Clostridium_sensu_stricto*, *Tepidimonas*, *Lysobacter* and *Lactobacillus* were present only in the Illumina dataset.

In general, there was a great overlap in the principal genera detected in both analyses, although with greater variability in relative abundances than those observed for ICU samples. As seen in **Figure 3A**, main genera in the NICU include *Bacillus* (on average, 40.5% of relative abundance estimated with nanopore and 28.8% with Illumina), *Capnocytophaga* (14.6% with nanopore and 22.8% with Illumina), *Delftia* (18.58% with nanopore and 7.49% with Illumina), *Neisseria* (14.3% nanopore with and 12.5% with Illumina), *Stenotrophomonas* (12.3% with nanopore and 9.29% with Illumina), *Haemophilus* (13.4% with nanopore and 7.6% with Illumina), *Staphylococcus* (7.96% with nanopore and 6.4% with Illumina) and *Pseudomonas* (6.12% with nanopore and 5.07% with Illumina). In one sample, *Serratia* was detected at a relative abundance of 46% in the nanopore dataset, while it appeared at 9.50% in the Illumina dataset.

**Figure 3.**
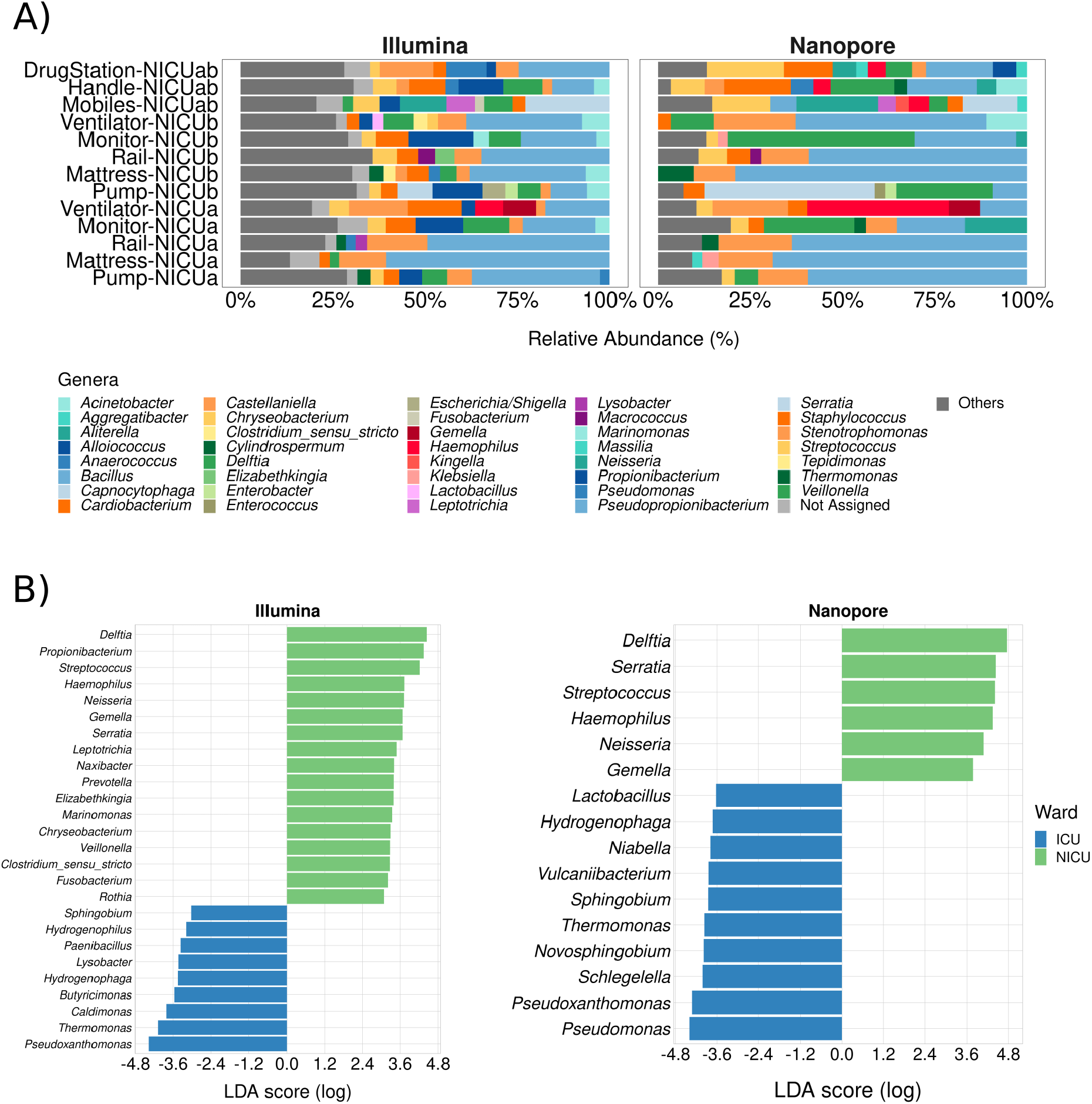
Neonatal Intensive Care Units microbial profiles obtained with Illumina and nanopore sequencing. **A)** Stacked bar plots depicting main genera detected in NICU samples using both methodologies. Genera with relative abundances below 2.5% were grouped under the label “Others”. **B)** Ward-specific **t**axonomic biomarkers for (N)ICU detected with linear discriminant analysis effect size (LEfSe). Statistical significance was defined as p < 0.05.

Both in our previous work and in the present study, NICU was shown to be more diverse than ICU **(Figure S1)** (9). With the 2.5% cutoff, an average of 26% of the genera present in NICU samples sequenced with Illumina fell under the label “Others”, highlighting the prevalence of low-abundance genera in these samples. In the nanopore dataset, on the other hand, low-abundance genera within the samples comprised, on average, only 10% of the total diversity, leading to a possible overestimation of the most abundant genera.

In order to assess whether these differences impact the determination of taxonomic biomarkers in the samples, we employed an algorithm for high-dimensional biomarker discovery that can associate genera specifically associated to the different wards (16). **Figure 3B** shows which genera were found to be associated with (N)ICU wards using both sequencing datasets. As expected, nanopore sequencing retrieved fewer site-specific genera when compared to Illumina, however, all of the genera associated with NICU with higher LDA scores (apart from *Propionibacterium*) were equally found in both datasets, while ICU taxonomic biomarkers showed more variation. As a whole, these results reinforce that nanopore’s capacity to detect less abundant genera could have been undermined by the stark contrast in throughput of each sequencing method.

### Full-length 16S rRNA sequencing provides insight into the species-level composition of ICU microbial communities

One great advantage of generating reads that encompass the entire length of the 16S rRNA gene is that taxonomic assignment can be performed by direct mapping of the query reads to the 16S reference database, which ultimately may associate each read to a particular bacterial species (18).

While nanopore methods may still lack accuracy to enable exact species-level correspondence between sequenced reads and organisms (19), a detailed look into the species assigned to our nanopore reads may show revealing features of the microbial communities and provide clues to how HAI-related organisms can be able to circumvent the cleaning procedures and underlying motions of cross-contamination between hospital areas. **Figure 4A** and **Figure 4B** show the top assigned species to each sample. While many genera are represented by more than one species, *Escherichia fergusonii, Pseudomonas thermotolerans. Pseudoproprionibacterium rubrum* and *K. pneumoniae* are examples of species found in elevated relative abundances that were often the only members of their genera assigned in samples.

**Figure 4.**
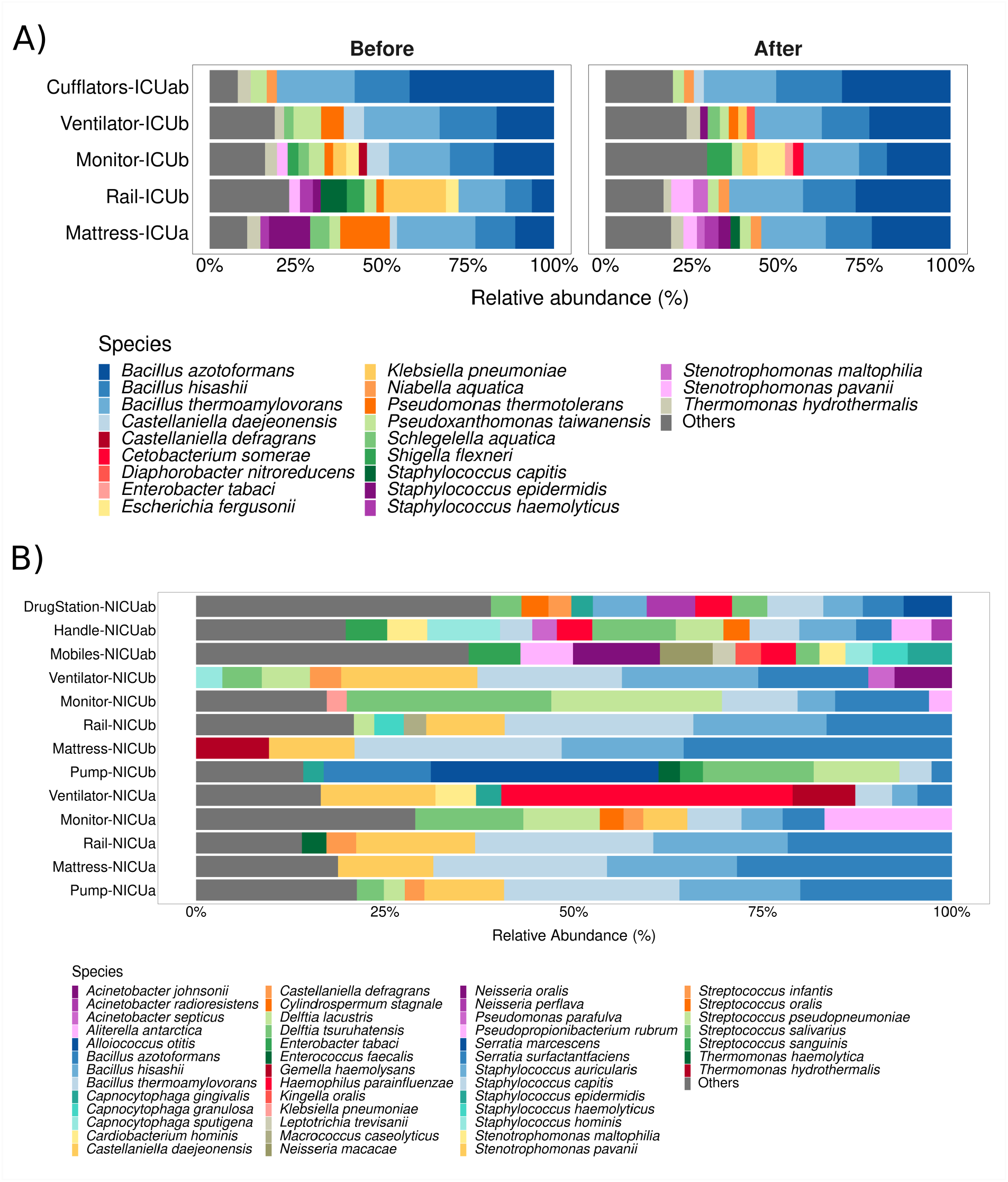
Species-level taxonomic assignment to (N)ICU samples. **A)** and **B)** depict the main species assigned to nanopore-generated reads for ICU and NICU samples, respectively. Species with relative abundances below 2% for ICU samples or 2.5% for NICU samples were grouped into the label “Others”.

## Discussion

Nanopore sequencing has emerged with the potential to radically change the landscape of the-omics sciences, and researchers are still becoming acquainted to the full breadth of possibilities it offers. As the community grows, standardization and validation of wet lab protocols and computational pipelines are imperative and, while efforts in this direction have been making great strides, there is still much room for improvement (6,20).

Amplification biases are a frequent concern during library preparation in massive parallel sequencing experiments, regardless of the sequencing platform employed (21). In this study, we could amplify only 32 out of 42 selected samples using Oxford Nanopore’s commercial set of primers and a standard PCR protocol, obtaining different degrees of success for each sample. Although it should be noted that the concentrations and qualities of our metagenomic templates were uneven, and that amplification could probably be achieved if reaction conditions were finetuned for each sample (which is ultimately impractical in a real-life setting), we believe that this is the most likely cause for the observed disparity in the number of reads obtained across samples.

Literature shows that the use of Oxford Nanopore’s commercial barcoding primers also potentially interfere with one’s capacity to detect certain bacterial taxa in complex samples. *Corynebacterium, Pseudomonas* and *Bifidobacterium* have been reported to be some of the affected genera (19,22,23). In our experiments, while we were able to detect *Pseudomonas* in relative abundances comparable to those previously observed in the Illumina experiments, *Corynebacterium* was indeed severely underrepresented. Moreover, we could not detect *Proprionibacterium* in our nanopore dataset, even though it was detected as a NICU biomarker by Illumina sequencing. Approaches such as the design of custom primers have been proved to be effective in circumventing amplification limitations and providing more accurate community profiles with nanopore sequencing efforts (23).

Aside from amplification issues, the low number of usable reads after processing steps was another striking result we observed. Literature suggests that this is not uncommon in nanopore sequencing studies. The relatively high error rate of current basecalling algorithms is a known limitation of the method, and researchers often have to define arbitrary quality thresholds in order to balance reliability of information with loss of data. Moreover, size range cutoffs chosen to filter reads certainly have an impact on the number of reads retrieved in a given experiment. Thus, most reports for microbial community analysis using 16S we found applied different Q Score thresholds (usually Q Score > 7) and variable cutoff values (e.g. 1300–1950bp, <1700bp, 1400-1700 bp, and 1350-1650 bp) (19,22–24).

Despite the variability in experiment design choices, there is a general consensus that nanopore sequencing is robust enough to provide reliable community composition information, at least at genus level. Our results are favorable to this conclusion. Even though the differences in throughput led to different diversity estimates between nanopore and Illumina datasets, the main genera detected in the samples were, with few noted exceptions, the same with both approaches and relative abundances were mostly comparable. Moreover, aside from assessing nanopore by comparison with Illumina, our analysis might have revealed possible shortcomings of the previous Illumina sequencing effort itself. Aside from the remarkable observation that the detection of *Klebsiella* was only possible with nanopore sequencing, even though this genus was found to be an important pathogen found in the hospital at that time, the absence of *Methylobacterium,* a known laboratory contaminant (25,26), in our nanopore dataset suggests that the high abundances of this genus previously reported in the sample monitor-ICUb was due to contamination during sample processing.

While promising, 16S-based species-level microbial profiling using nanopore remains a rather controversial topic. Current error rates reported for long-read basecalling algorithms are usually higher than the differences between closely related species, which makes species-level assignment unadvisable due to the possibility of false positives (19,22). Using a mock community, Winand and collaborators reported that, even though genus-level classification of nanopore is highly accurate, the use of absolute number of reads as a “tiebreak” between organisms of the same genus would not reflect the real composition of the community for all present species (19). Other works are more favorable to species-level taxonomic assignment using nanopore, arguing that low discrimination of closely related species (e.g., members of the genera *Bacillus* and *Escherichia)* is a limitation of using 16S as a biomarker by itself, and could be overcome if more comprehensive sequences, such as the entire *rrn* operon (16S rRNA-ITS-23S rRNA), were employed instead (18,23).

Upon investigation, we found that species-level taxonomic assignment to our nanopore reads indeed resulted in split categorization between different species within the represented genera. Reads belonging to the genus *Bacillus,* for instance, were assigned to *Bacillus azotoformans, Bacillus hisahii* and *Bacillus thermoamylovorans* in most samples. Without any additional information, it would be impossible to determine which of these species were actually present in the environments and their actual relative abundances. However, we believe that some of the results, although inconclusive, may direct further investigation. For instance, it has caught our attention that the most abundant *Escherichia* species in the ICU samples was *E. fergusonii.* In fact, in many samples it has been the sole *Escherichia* detected. While still considered a novel pathogen, a recent publication reported the presence of extended-spectrum β-lactamase-producing *E. fergusonii* in poultry farms in the state of São Paulo (27), which stresses the importance of monitoring this microorganism in healthcare facilities of our region. In the NICU samples, the knowledge that *Haemophilu sparainfluenzae* is the sole representative of its genera in samples like ventilators might be an important clue to support accurate diagnostics of *Haemophilus* spp. nosocomial infections, since traditional phenotypic assays can lead to faulty results (28).

Nonetheless, hospital environments such as NICUs are known to house several related species concurrently, and finding ways to accurately detect different related bacteria at species- or even strain-level is important for effective surveillance and diagnosis in healthcare settings (29,30). Recently developed tools like the Bonito basecaller promise to take on some of nanopore’s main limitations and may turn high-accuracy long-read sequencing into a reality within the next years (31).

To us, it is exciting that a small team and a couple of days worth of bench work were enough to verify findings that were previously obtained by a much larger cohort of our colleagues, all without the cumbersome requirement of sending samples back and forth to special facilities. As methods become more refined with time, nanopore sequencing certainly has the potential to empower healthcare settings to monitor environmental threats rapidly and reliably on a day-to-day basis, revolutionizing healthcare as we know it today.

## Author Contributions

M-EG and RS-R contributed to conception and design of the study. GMVS performed the wet-lab procedures, including 16S amplification and nanopore sequencing. GMVS and FMP-d-S wrote the scripts for processing raw long-read data. GMVS conducted the data analysis and wrote the first draft of the manuscript. All authors contributed to manuscript revision, read, and approved the submitted version

## Acknowledgments

The authors would like to thank Lucas Ribeiro for his work in collecting samples and prominent contribution to the original publication, and all the colleagues of our group for their insightful comments in the course of this work.

## Supplementary Material

The Supplementary Material for this article can be found at the end of this document

## Data Availability Statement

The nucleotide sequences obtained in the present study have been deposited in the Sequenced Read Archive (SRA) database under the Accession number PRJNA728896.

## References

1. Weber DJ, Anderson D, Rutala WA. The role of the surface environment in healthcare-associated infections. Curr Opin Infect Dis. 2013;26(4):338–44.

2. Suleyman G, Alangaden G, Bardossy AC. The Role of Environmental Contamination in the Transmission of Nosocomial Pathogens and Healthcare-Associated Infections. Curr Infect Dis Rep. 2018;20(6).

3. Comar M, D’accolti M, Cason C, Soffritti I, Campisciano G, Lanzoni L, et al. Introduction of NGS in environmental surveillance for healthcare-associated infection control. Microorganisms. 2019;7(12).

4. Váradi L, Luo JL, Hibbs DE, Perry JD, Anderson RJ, Orenga S, et al. Methods for the detection and identification of pathogenic bacteria: Past, present, and future. Chem Soc Rev [Internet]. 2017;46(16):4818–32. Available from: http://dx.doi.org/10.1039/C6CS00693K

5. Jain M, Olsen HE, Paten B, Akeson M. The Oxford Nanopore MinION: delivery of nanopore sequencing to the genomics community. Genome Biol [Internet]. 2016;17(1):1–11. Available from: http://dx.doi.org/10.1186/s13059-016-1103-0

6. Ciuffreda L, Rodríguez-Pérez H, Flores C. Nanopore sequencing and its application to the study of microbial communities. Comput Struct Biotechnol J [Internet]. 2021;19:1497–511. Available from: https://doi.org/10.1016/j.csbj.2021.02.020

7. Tremblay J, Singh K, Fern A, Kirton ES, He S, Woyke T, et al. Primer and platform effects on 16S rRNA tag sequencing. Front Microbiol. 2015;6(AUG):1–15.

8. Petersen LM, Martin IW, Moschetti WE, Kershaw CM, Tsongalis GJ. Third-Generation Sequencing in the Clinical Laboratory: Sequencing. J Clin Microbiol. 2019;58(1):1–10.

9. Ribeiro LF, Lopes EM, Kishi LT, Ribeiro LFC, Menegueti MG, Gaspar GG, et al. Microbial community profiling in intensive care units expose limitations in current sanitary standards. Front Public Heal. 2019;7(AUG):1–14.

10. De Coster W, D’Hert S, Schultz DT, Cruts M, Van Broeckhoven C. NanoPack: Visualizing and processing long-read sequencing data. Bioinformatics. 2018;34(15):2666–9.

11. Li H. Minimap2: Pairwise alignment for nucleotide sequences. Bioinformatics. 2018;34(18):3094–100.

12. Winter D, Lee K, Cox M. Package ‘ pafr.’ 2020;

13. Moritz P, Nishihara R, Wang S, Tumanov A, Liaw R, Liang E, et al. Ray: A distributed framework for emerging AI applications. Proc 13th USENIX Symp Oper Syst Des Implementation, OSDI 2018. 2007;561–77.

14. Oksanen AJ, Blanchet FG, Friendly M, Kindt R, Legendre P, Mcglinn D, et al. Package ‘ vegan.’ 2020;

15. Wickham H, Averick M, Bryan J, Chang W, McGowan L, François R, et al. Welcome to the Tidyverse. J Open Source Softw. 2019;4(43):1686.

16. Segata N, Izard J, Waldron L, Gevers D, Miropolsky L, Garrett WS, et al. Metagenomic biomarker discovery and explanation. Genome Biol [Internet]. 2011;12(6):R60. Available from: http://genomebiology.biomedcentral.com/articles/10.1186/gb-2011-12-6-r60

17. Devanga Ragupathi NK, Muthuirulandi Sethuvel DP, Inbanathan FY, Veeraraghavan B. Accurate differentiation of Escherichia coli and Shigella serogroups: challenges and strategies. New Microbes New Infect [Internet]. 2018;21:58–62. Available from: https://doi.org/10.1016/j.nmni.2017.09.003

18. Cuscó A, Catozzi C, Viñes J, Sanchez A, Francino O. Microbiota profiling with long amplicons using nanopore sequencing: Full-length 16s rRNA gene and whole rrn operon [version 1; referees: 2 approved, 3 approved with reservations]. F1000Research. 2018;7:1–29.

19. Winand R, Bogaerts B, Hoffman S, Lefevre L, Delvoye M, Van Braekel J, et al. Targeting the 16s rRNA gene for bacterial identification in complex mixed samples: Comparative evaluation of second (illumina) and third (oxford nanopore technologies) generation sequencing technologies. Int J Mol Sci. 2020;21(1):1–22.

20. Santos A, van Aerle R, Barrientos L, Martinez-Urtaza J. Computational methods for 16S metabarcoding studies using Nanopore sequencing data. Comput Struct Biotechnol J [Internet]. 2020;18:296–305. Available from: https://doi.org/10.1016/j.csbj.2020.01.005

21. Van Dijk EL, Jaszczyszyn Y, Thermes C. Library preparation methods for next-generation sequencing: Tone down the bias. Exp Cell Res [Internet]. 2014;322(1):12–20. Available from: http://dx.doi.org/10.1016/j.yexcr.2014.01.008

22. Heikema AP, Horst-Kreft D, Boers SA, Jansen R, Hiltemann SD, de Koning W, et al. Comparison of illumina versus nanopore 16s rRNA gene sequencing of the human nasal microbiota. Genes (Basel). 2020;11(9):1–17.

23. Matsuo Y, Komiya S, Yasumizu Y, Yasuoka Y, Mizushima K, Takagi T, et al. Full-length 16S rRNA gene amplicon analysis of human gut microbiota using MinION^™^ nanopore sequencing confers species-level resolution. BMC Microbiol. 2021;21(1):1–13.

24. Stahl-rommel S, Jain M, Nguyen HN, Arnold RR, Aunon-chancellor SM, Sharp GM, et al. the International Space Station Using Nanopore Sequencing. 2021;

25. Glassing A, Dowd SE, Galandiuk S, Davis B, Chiodini RJ. Inherent bacterial DNA contamination of extraction and sequencing reagents may affect interpretation of microbiota in low bacterial biomass samples. Gut Pathog. 2016;8(1):1–12.

26. Salter SJ, Cox MJ, Turek EM, Calus ST, Cookson WO, Moffatt MF, et al. Reagent and laboratory contamination can critically impact sequence-based microbiome analyses. BMC Biol. 2014;12(1):1–12.

27. Ferreira JC, Penha Filho RAC, Andrade LN, Berchieri Junior A, Darini ALC. Evaluation and characterization of plasmids carrying CTX-M genes in a non-clonal population of multidrug-resistant Enterobacteriaceae isolated from poultry in Brazil. Diagn Microbiol Infect Dis [Internet]. 2016;85(4):444–8. Available from: http://dx.doi.org/10.1016/j.diagmicrobio.2016.05.011

28. Andrzejczuk S, Kosikowska U, Chwiejczak E, Stępień-Pyśniak D, Malm A. Prevalence of resistance to β-Lactam antibiotics and bla genes among commensal Haemophilus parainfluenzae isolates from respiratory microbiota in Poland. Microorganisms. 2019;7(10).

29. Tett A, Pasolli E, Farina S, Truong DT, Asnicar F, Zolfo M, et al. Unexplored diversity and strain-level structure of the skin microbiome associated with psoriasis. npj Biofilms Microbiomes [Internet]. 2017;3(1):1–11. Available from: http://dx.doi.org/10.1038/s41522-017-0022-5

30. Brooks B, Olm MR, Firek BA, Baker R, Thomas BC, Morowitz MJ, et al. Strain-resolved analysis of hospital rooms and infants reveals overlap between the human and room microbiome. Nat Commun. 2017;8(1):1–7.

31. Vereecke N, Bokma J, Haesebrouck F, Nauwynck H, Boyen F, Pardon B, et al. High quality genome assemblies of Mycoplasma bovis using a taxon-specific Bonito basecaller for MinION and Flongle long-read nanopore sequencing. BMC Bioinformatics [Internet]. 2020;21(1):1–16. Available from: https://doi.org/10.1186/s12859-020-03856-0

