## Supplementary Material, table S1, figures S1, S2 and S3. for "Nanopore sequencing provides rapid and reliable insight into microbial profiles of Intensive Care Units"

**Table S1.** Summary statistics of nanopore sequencing data for (N)ICU surface samples

| | after basecalling | | | | after filtering and QC | | | |
|---|---|---|---|---|---|---|---|---|
| Sample | max (bp) | min (bp) | avg (bp) | total reads | max (bp) | min (bp) | avg (bp) | total reads |
| Card-ICUab | 4226 | 36 | 1450.709 | 17317 | 1569 | 1351 | 1439.334 | 1418 |
| Cufflators-ICUab | 4081 | 51 | 1451.937 | 7174 | 1559 | 1355 | 1442.184 | 565 |
| Cufflators-ICUabA | 3159 | 71 | 1446.591 | 10538 | 1621 | 1352 | 1448.104 | 2944 |
| DrugStation-ICUab | 3140 | 75 | 1460.719 | 2152 | 1534 | 1352 | 1446.408 | 174 |
| DrugStation-NICUab | 3112 | 59 | 1444.486 | 6102 | 1597 | 1355 | 1444.446 | 1750 |
| Handle-ICUab | 3217 | 34 | 1435.297 | 14752 | 1579 | 1351 | 1434.405 | 1222 |
| Handle-NICUab | 3024 | 53 | 1436.484 | 2882 | 1565 | 1350 | 1439.511 | 824 |
| Mattress-ICUa | 3273 | 32 | 1450.524 | 12027 | 1619 | 1356 | 1442.259 | 1039 |
| Mattress-ICUaA | 3865 | 10 | 1431.443 | 31276 | 1581 | 1350 | 1448.602 | 8585 |
| Mattress-ICUb | 3119 | 41 | 1440.604 | 15215 | 1646 | 1351 | 1446.253 | 4551 |
| Mattress-NICUa | 3089 | 81 | 1446.941 | 8863 | 1638 | 1350 | 1448.1 | 2581 |
| Mattress-NICUb | 3128 | 50 | 1464.799 | 1635 | 1602 | 1353 | 1445.525 | 139 |
| MedicalRecord-ICUab | 3203 | 44 | 1402.943 | 19358 | 1626 | 1350 | 1436.101 | 1520 |
| Mobiles-NICUab | 3214 | 52 | 1434.405 | 21184 | 1629 | 1350 | 1440.422 | 6125 |
| Monitor-ICUaA | 3158 | 66 | 1444.956 | 14770 | 1637 | 1354 | 1444.867 | 4019 |
| Monitor-ICUb | 3106 | 53 | 1440.017 | 8925 | 1595 | 1350 | 1443.119 | 2655 |
| Monitor-ICUbA | 3308 | 36 | 1405.712 | 27027 | 1608 | 1350 | 1439.851 | 2266 |
| Monitor-NICUa | 15467 | 16 | 1427.924 | 14335 | 1595 | 1350 | 1432.931 | 4200 |
| Monitor-NICUb | 3207 | 30 | 1379.87 | 13543 | 1581 | 1350 | 1434.558 | 1106 |
| Pump-ICUa | 4113 | 24 | 1391.855 | 34083 | 1587 | 1350 | 1441.986 | 2541 |
| Pump-ICUb | 3182 | 10 | 1451.5 | 11899 | 1601 | 1354 | 1453.717 | 3627 |
| Pump-NICUa | 3211 | 54 | 1417.056 | 16759 | 1617 | 1350 | 1446.297 | 4572 |
| Pump-NICUb | 3174 | 59 | 1440.717 | 8088 | 1553 | 1353 | 1439.45 | 754 |
| Rail-ICUb | 3227 | 55 | 1423.863 | 16891 | 1598 | 1351 | 1442.948 | 4895 |
| Rail-ICUbA | 3251 | 36 | 1430.81 | 27337 | 1590 | 1350 | 1448.805 | 2253 |
| Rail-NICUa | 3195 | 58 | 1417.085 | 17144 | 1616 | 1350 | 1445.584 | 4718 |
| Rail-NICUb | 4720 | 24 | 1397.072 | 19717 | 1610 | 1354 | 1441.449 | 1513 |
| Ventilator-ICUaA | 3158 | 80 | 1448.31 | 1611 | 1588 | 1376 | 1448.026 | 461 |
| Ventilator-ICUb | 3147 | 52 | 1443.374 | 10911 | 1626 | 1351 | 1446.121 | 3245 |
| Ventilator-ICUbA | 3393 | 24 | 1408.932 | 25273 | 1575 | 1352 | 1437.071 | 1914 |
| Ventilator-NICUa | 3200 | 54 | 1442.009 | 14947 | 1604 | 1350 | 1449.432 | 4148 |
| Ventilator-NICUb | 4782 | 43 | 1454.293 | 3735 | 1566 | 1355 | 1439.678 | 298 |

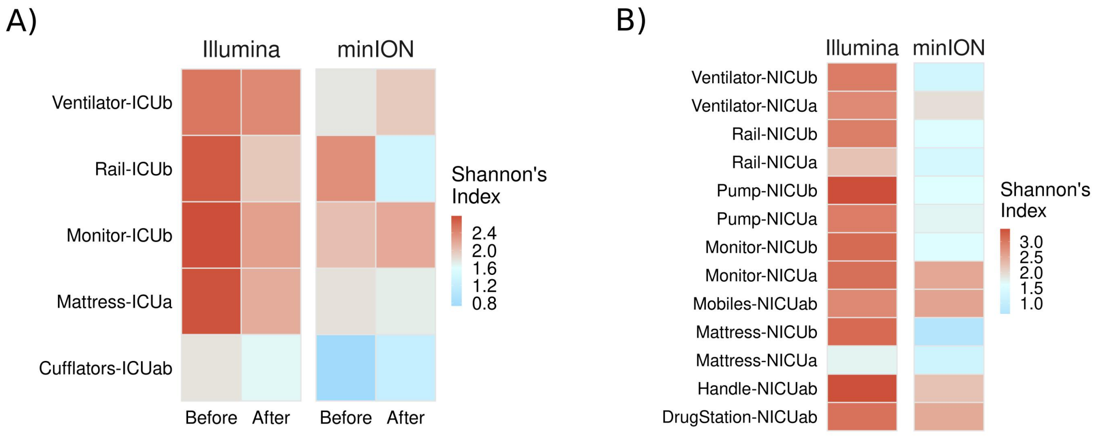


**Figure S1 -** Alpha diversity (Shannon Index) at genus level measured for samples from the nanopore and Illumina datasets. **A)** ICU samples before and after cleaning and **B)** NICU samples.

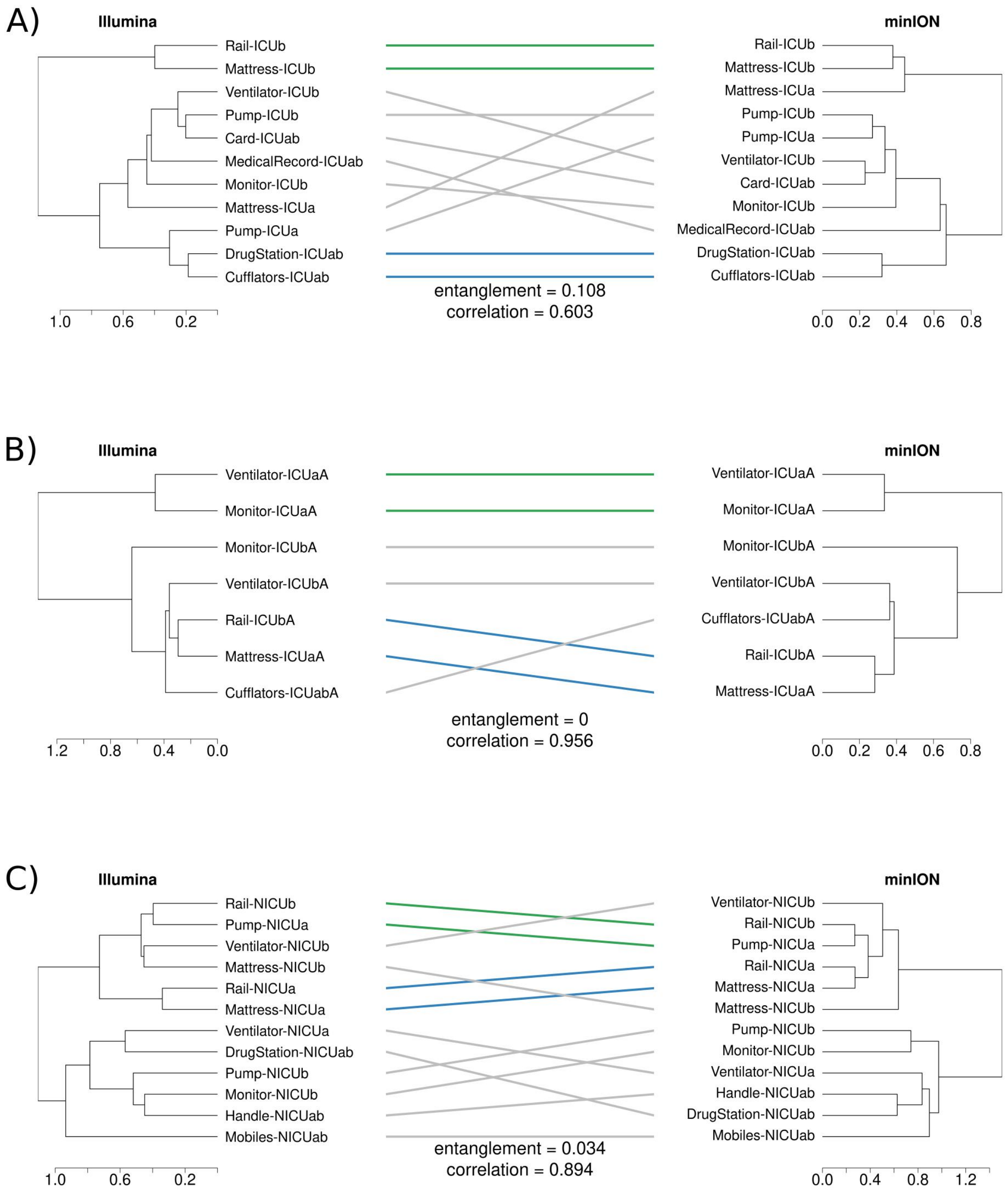


**Figure S2 -** Tanglegrams showing the hierarchical clustering (Ward's algorithm) similarities between samples sequenced with Illumina and nanopore in **A)** ICU samples before the concurrent cleaning, **B)** ICU samples after the concurrent cleaning, and **C)** NICU samples. Distance between groups was measured using Jaccard index. Correlation and entanglement methods employed were, respectively cophenetic and "step2side". Horizontal rulers indicate the clustering heights.

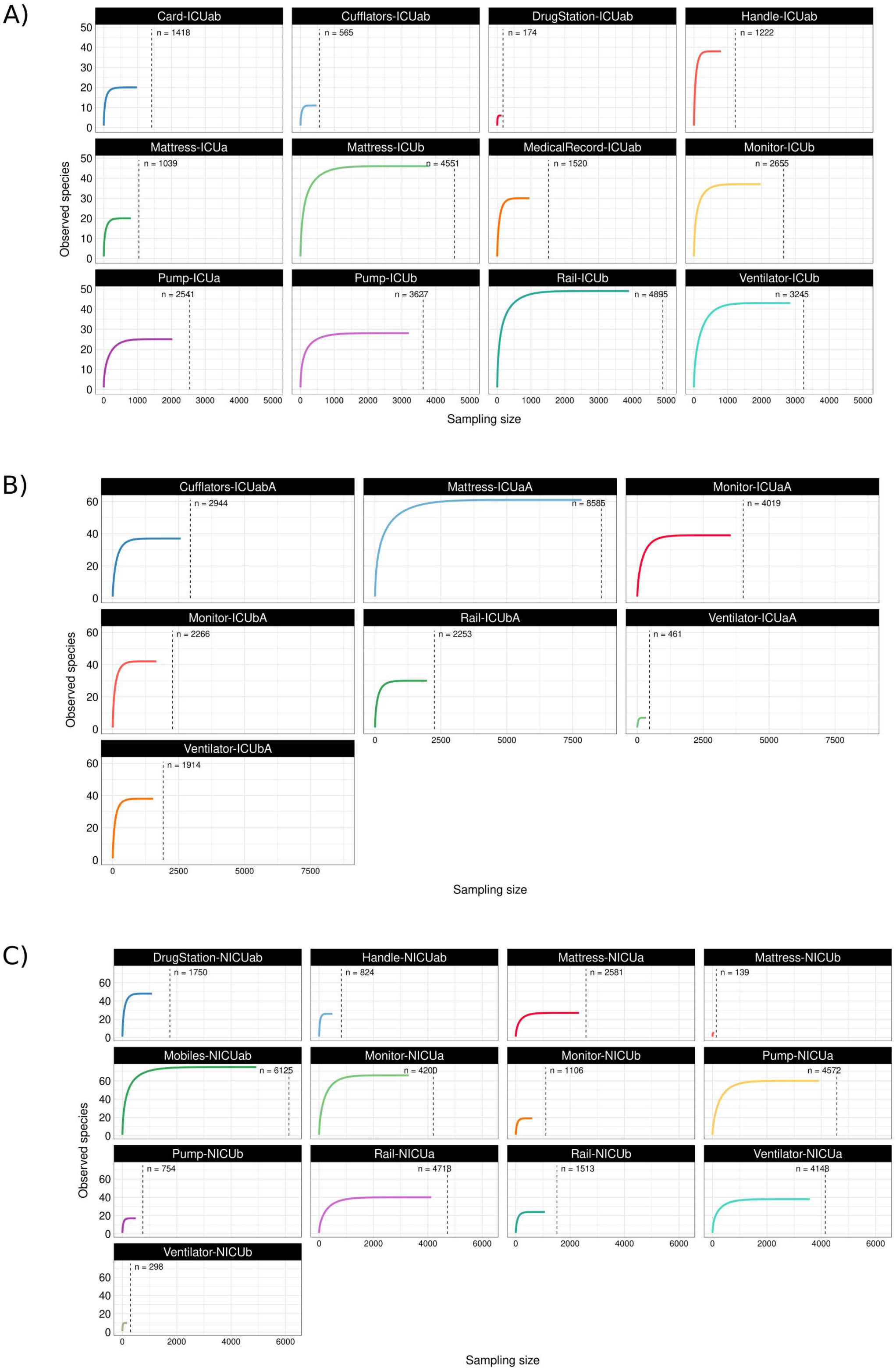


**Figure S3 -** Rarefaction curves for nanopore-sequenced samples. **A)** samples from ICU before the cleaning, **B)** samples from ICU after cleaning, **C)** samples from NICU. The dashed line demarcates the *n* number of reads that remained after all processing steps.
